# Cereblon promotes influenza virus replication through AMPK ubiquitination

**DOI:** 10.64898/2025.12.17.694861

**Authors:** Thu Ha Nguyen, Dong Ju Lee, Mahmoud Soliman, Muhammad Sharif, Myra Hosmillo, Hyung-Jun Kwon, Seong-Hun Jeong, Dae-Eun Cheong, Hae-Rang Seo, Sunwoo Lee, Don-Kyu Kim, Chul-Seung Park, Hueng-Sik Choi, Tae-Il Jeon, Ian G. Goodfellow, Kyoung-Oh Cho

## Abstract

Influenza A and B viruses (IAV and IBV) continually threaten global health, with IAV posing a risk of emerging pandemics. Rapid viral evolution makes current treatments less effective, highlighting the urgent need for broad-spectrum antivirals. Targeting host factors essential for viral replication may offer a highly promising broad-spectrum antiviral strategy. In this context, cereblon (CRBN), a substrate adaptor of the CRL4 E3 ubiquitin ligase complex, is found to promote both IAV and IBV replication as a key pro-viral host factor. Mechanistically, CRBN targets and degrades AMP-activated protein kinase (AMPK) via the proteasome. This CRBN-driven degradation shifts the metabolism of the infected cells toward anabolism, promoting lipid droplet (LD) formation and creating a microenvironment favorable for viral replication. Genetic depletion or inhibition of CRBN stabilizes AMPK, significantly reduces LD formation, and effectively suppresses the replication of various IAV and IBV strains in vivo, demonstrating its broad-spectrum potential. Notably, Crbn knockout mice show marked resistance to lethal IAV infection. CRBN inhibition with immunomodulatory imide drugs—known CRBN inhibitors—significantly decreases IAV replication in vivo. This research underscores CRBN as a crucial regulator of host metabolism during viral infection, revealing the CRBN-AMPK axis as a promising target for host-directed pan-influenza antiviral development.

**Author Summary:** Influenza A and B viruses are response for recurring seasonal epidemics and occasional pandemics that pose serious global health challenges. Current antiviral drugs often lose effectiveness as influenza viruses rapidly develop resistance through genetic mutations. To overcome this limitation, our study focused on a host factor targeted by influenza viruses-the ubiquitin E3 ligase substrate adaptor cereblon (CRBN). We discovered that influenza viruses exploit CRBN to enhance their replication by altering host cell metabolism. Specifically, CRBN binds to a key energy sensor – AMPKα and induces the ubiquitination and degradation of AMPKγ. Consequently, the AMPK suppression shifts the cellular environment from catabolism to anabolism, activating lipogenic enzymes and promoting lipid droplets formation which provide a favorable environment for viral growth. Our study highlight that blocking CRBN either genetically or chemical inhibition (immunomodulatory imide drugs) of CRBN reduces LD formation and strongly suppresses diverse influenza strains replication in vitro and in vivo. These findings reveal not only how viruses hijack host metabolism via the CRBN-AMPK pathway but also present a potential therapeutic for novel broad-spectrum anti-influenza.

## Introduction

Influenza A and B viruses (IAV and IBV) cause seasonal flu outbreaks worldwide, leading to approximately 290,000 to 650,000 deaths each year [1, 2]. IAVs have also been linked to severe pandemics in human history, such as the H1N1 Spanish influenza outbreak in 1918, the H2N2 Asian influenza outbreak in 1957, the H3N2 Hong Kong influenza outbreak in 1968, and the H1N1 swine influenza outbreak in 2009 [3, 4]. The recent global surge of highly pathogenic avian IAV H5N1 spreading into various domestic and wild animals highlights the ongoing pandemic threat posed by IAVs [4–7]. The persistent threat and rapid evolution of IAVs underscore the need for developing vaccines and therapeutics that are effective not only against currently circulating strains but also against any emerging new ones. Currently available antivirals against influenza may become less effective or even ineffective against mutant or emerging IAVs [8, 9]. Therefore, there is a continuing need to develop next-generation antivirals, especially those targeting host factors essential for the virus’s life cycle, as they are likely to offer a higher barrier to resistance than direct-acting therapeutics [10–13].

The dynamic interplay between viruses and host cells reshapes the intracellular microenvironment by driving metabolic reprogramming to support viral replication [14–16]. In particular, viruses hijack key metabolic pathways, such as fatty acid biosynthesis, glycolysis, and glutaminolysis, to optimize their lifecycle [14–16]. Viral infection and the resulting host responses can compromise the integrity of cellular membranes [17–19] disrupting nutrient uptake and negatively impacting viral replication [17, 20, 21]. To counter this, viruses often promote the buildup of lipid droplets (LDs) early during infection, ensuring a continuous supply of essential lipids [17, 22]. LDs serve as reservoirs for free fatty acids (FFAs), which contribute to the formation of viral replication compartments for positive-sense RNA viruses, fuel energy production through mitochondrial fatty acid oxidation, and facilitate the palmitoylation of viral proteins [17, 22–25].

Cereblon (CRBN), a substrate receptor of the E3 ubiquitin ligase complex, ubiquitinates numerous intracellular proteins, including a component of the AMP-activated protein kinase (AMPK) complex [26–31]. As AMPK switches on ATP-generating catabolic pathways, CRBN-mediated degradation of AMPK enhances anabolic processes such as fatty acid biosynthesis [27, 29, 32, 33]. Conversely, inhibiting CRBN activates AMPK, shifting the metabolism towards catabolic energy production and preventing lipid accumulation, as seen in high-fat diet *Crbn*-knockout (KO) mice [33].

In this study, we demonstrate that virus-induced CRBN activation promotes ubiquitination and degradation of AMPK complexes, reshaping the intracellular microenvironment and driving LD formation, a process essential for virus replication [17, 22–25]. Chemical inhibition or knockdown of CRBN in virus-infected cells is shown to restore AMPK activity, reducing LD formation and impairing viral progeny production. Compared to wild-type (WT) mice, *Crbn* KO mice exhibit resistance to IAV-induced mortality, and chemical inhibition of CRBN conferred protection against lethal IAV infection. These findings provide new insights into the role of CRBN-mediated AMPK regulation in viral infection. These findings align with recent observations that a deficiency in CRBN reduces IAV infection both *in vitro* and *in vivo* [34] further advancing our understanding of the role of CRBN-mediated AMPK function in viral infection and potentially facilitating the rational design of safer antiviral therapeutics targeting the CRBN-AMPK axis.

## Results

### CRBN Deficiency Confers Resistance to IAV Infection

LDs are crucial for the replication of many viruses [17, 22–25, 35–39]. Previously, we demonstrated that Crbn knockout suppresses hepatic LD accumulation under high-fat conditions [33] potentially indicating that CRBN activity may in some way be linked to LD-mediated viral replication efficiency. To explore the role of CRBN in virus replication, we assessed the susceptibility of Crbn KO mice to IAV challenge. The genotypes of the WT (Crbn^+/+^), heterozygous KO (Crbn^+/-^), and homozygous KO (Crbn^-/-^) C57BL/6N mice were confirmed through PCR Fig 1A and western blot analysis Fig 1B [33]. All WT mice challenged with the mouse-adapted PR8 strain died within 10 days, while 80% of the homozygous KO (Crbn^-/-^) mice survived after the IAV challenge Fig 1C. Body weight losses and clinical scores following IAV infection were significantly reduced in Crbn KO mice compared to WT mice Fig 1D-E. Viral protein expression and genome copy numbers were markedly lower in the lung samples from IAV-challenged Crbn KO mice than in WT mice Fig 1F-G. Histopathological and immunohistochemical analyses revealed that IAV-induced lung lesions and distribution of IAV-positive cells in the WT mice, characterized by infiltration of macrophages, lymphocytes, and neutrophils in the interstitium and alveolar lumen and diffuse IAV-positive cells in the bronchiolar epithelial cells and alveolar pneumocytes, were significantly alleviated in Crbn KO mice Fig 1H. Collectively, these findings suggest that CRBN may, in some way, support IAV replication in vivo.

**Figure 1.**
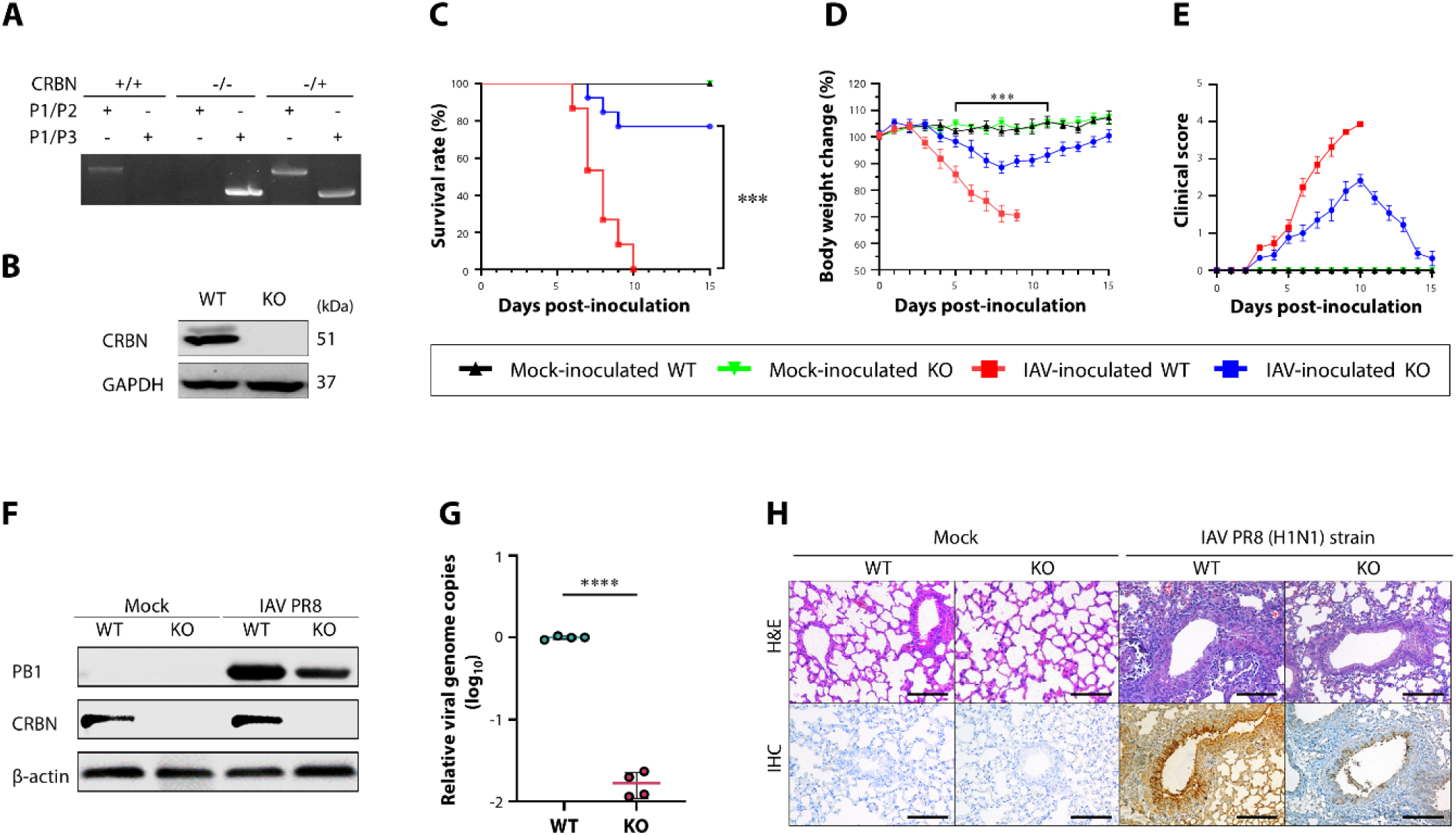
CRBN knockout (KO) mice are resistant to influenza A virus infection. (A) Representative image of PCR analysis of the *Crbn* gene from the tails of the wild type (WT, *Crbn^+/+^*), heterozygous KO (*Crbn^+/-^*), and homozygous KO (*Crbn^-/-^*) C57BL/6N mice using primer pair P1 and P2 and primer pair P1 and P3. (B) Representative images of immunoblot analysis of CRBN expression levels in the tails from WT and *Crbn^-/-^* mice. GAPDH was used as a loading control. (C-E) Comparison of survival rates (C), body weight changes (D), and clinical scores (E) between WT and homozygous KO (*Crbn^-/-^*) mice after challenge with 10^3^ PFU of mouse-adapted IAV PR8 strain (n = 16). (F and G) Comparison of IAV PB1 protein (F) and genome copy numbers (G) in the lungs from WT and homozygous KO (*Crbn^-/-^*) mice after challenge with 10^3^ PFU of mouse-adapted IAV PR8 strain. β-actin was used as a loading control. (H) Representative images of histological lesion changes (upper panels) and immunohistochemical changes (lower panels) of viral antigen distribution in the lungs of WT and CRBN KO mice days after challenge with or without the 10^3^ PFU of mouse-adapted IAV PR8 strain. Hematoxylin and eosin stain. Immunohistochemistry with an antibody against the IAV virion protein. Results are presented as arithmetic means ± S.D. Differences were evaluated using a one-way ANOVA. \**p* < 0.05; \*\**p* < 0.001; \*\*\**p* < 0.0001; \*\*\*\**p* < 0.00001. The scale bars in panel H correspond to 800 µm.

### CRBN-Dependent Modulation of AMPK and LD Dynamics during IAV Infection

Given the established correlation between CRBN activity and LD accumulation [33] we investigated how CRBN interconnects with LD dynamics during viral infection. Consistent with our previous findings [17] we observed that LD numbers in IAV-infected cells increased from the early stage of infection, peaked at the mid-stage, and subsequently decreased Fig 2A. Notably, CRBN expression followed a similar temporal pattern, with both mRNA and protein levels in response to IAV infection in vitro Fig 2B-D and in lung tissues from IAV-infected mice Fig 2E-H.

**Figure 2.**
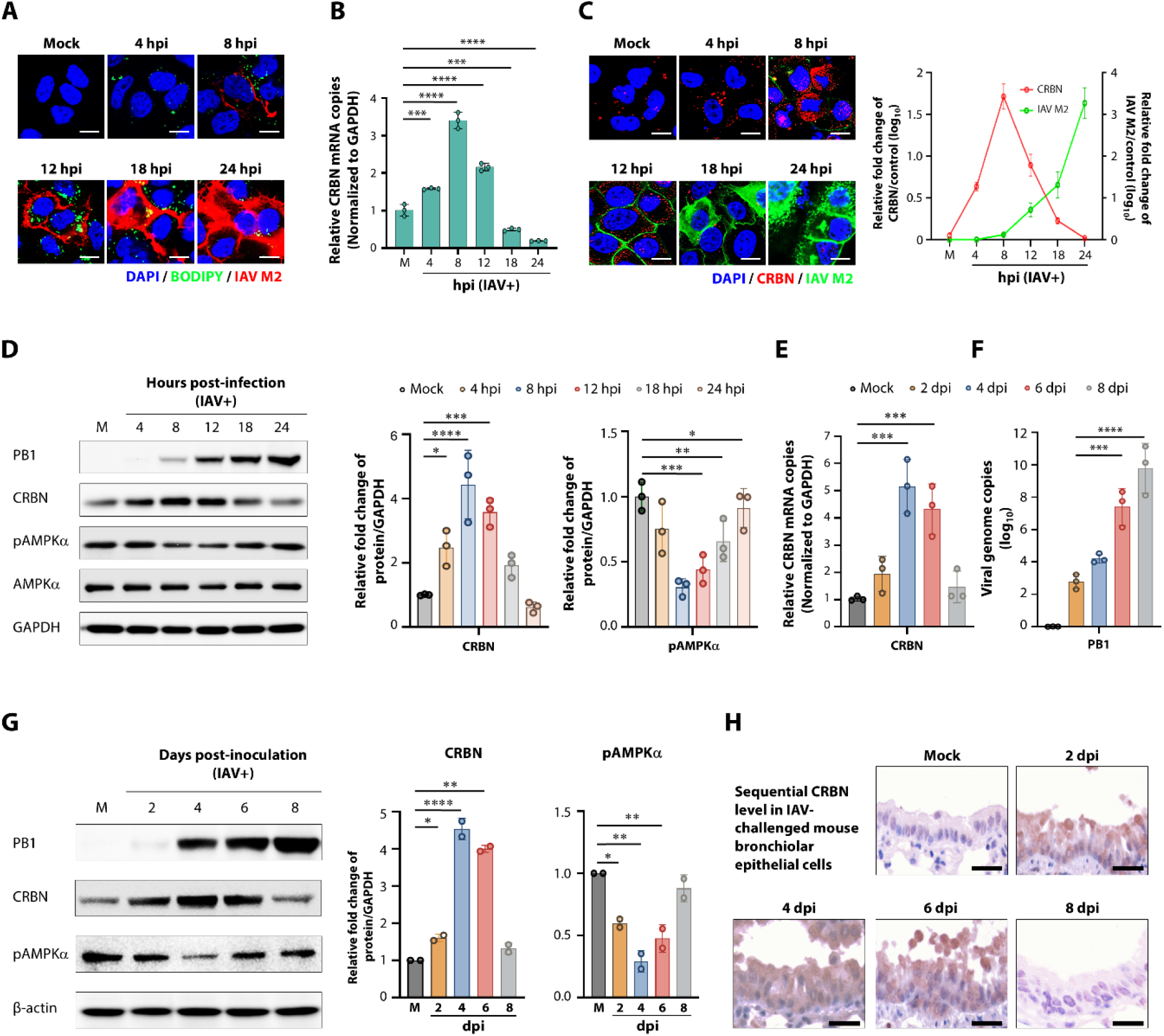
CRBN dynamic is similar to LD one but opposite to AMPK one. (A) Representative images of sequential changes of BODIPY-stained intracellular LDs (green) and IAV M2 protein (red) in A549 cells infected with IAV PR8 strain at an MOI of 1.0 FFU. (B) Sequential quantification of transcription level of the *Crbn* gene in A549 cells infected with the IAV PR8 strain at an MOI of 1 FFU by RT-qPCR analysis. (C) Representative images (left) and graphical representation (right) of sequential changes of CRBN (red) and IAV M2 protein (green) in A549 cells infected with IAV PR8 strain at an MOI of 1.0 FFU. (D) Representative western blot images (left) and quantification (right) of sequential changes of CRBN and pAMPKα in A549 cells infected with IAV PR8 strain at an MOI of 1.0 FFU. GAPDH was used as a loading control. (E-F) Quantification of sequential changes of mRNA of *Crbn* gene (E) and IAV genome copy numbers (F) in the mice challenged with 10^3^ PFU of mouse-adapted IAV PR8 strain by RT-qPCR. (G) Representative western blot images (left) and graphical representation (right) of sequential changes of CRBN and pAMPKα p in the lung samples from the mice challenged with 10^3^ PFU of mouse-adapted IAV PR8 strain. (H) Representative images of sequential changes of CRBN levels in the bronchiolar epithelial cells from the mice challenged with 10^3^ PFU of the mouse-adapted IAV PR8 strain. Immunohistochemistry with an antibody against CRBN. Results are presented as arithmetic means ± S.D. Differences were evaluated using a one-way ANOVA. \**p* < 0.05; \*\**p* < 0.001; \*\*\**p* < 0.0001; \*\*\*\**p* < 0.00001. The scale bars in panels A, C, and H correspond to 10 µm, 10 µm, and 100 µm, respectively.

CRBN has been shown to promote AMPK degradation via the ubiquitin-proteasome system, thereby shifting cellular metabolism toward anabolic pathways and facilitating LD formation [32] Based on this, we hypothesized that the dynamics of phosphorylated AMPKα (pAMPKα) would inversely correlate with CRBN expression and LD formation during viral infection. Notably, pAMPKα levels decreased during the early and mid-stages of IAV infection, which was inversely correlated with the increase in CRBN and LD in both IAV-infected cells and mouse lung tissues Fig 2D and G. To determine whether this CRBN-AMPK axis extends to other viruses that rely on LDs for replication, we examined cells infected with bovine coronavirus (BCoV), porcine epidemic diarrhea coronavirus (PEDV), bovine species A rotavirus (RVA), porcine reproductive and respiratory syndrome virus (PRRSV), or porcine sapovirus (PSaV). Across all tested viruses, we observed similar CRBN induction and concurrent reductions in pAMPKα levels during infection S1 Fig. These results suggest that CRBN ubiquitinates AMPK complexes to reprogram host cell metabolism, facilitating LD accumulation and supporting the replication of IAV and other LD-dependent viruses.

### Direct Interaction between CRBN and AMPK during Virus Replication

To determine if CRBN directly targets AMPKα during viral replication, we first analyzed global ubiquitination levels in cells infected with IAV. Ubiquitinated proteins increased for up to 8 hours post-infection (hpi) before declining, a pattern that paralleled CRBN expression but was inversely correlated with pAMPKα levels Fig 3A. We next tested whether AMPKα is a direct substrate of CRBN-mediated ubiquitination. Immunoprecipitation assays confirmed a physical interaction between CRBN and AMPKα, with reciprocal co-immunoprecipitation observed Fig 3B and S2A Fig. Notably, the binding dynamics of CRBN and pAMPKα mirrored CRBN expression levels and were inversely related to pAMPKα abundance. The interaction between CRBN and AMPKα was further validated using a Duolink proximity ligation assay (PLA), which demonstrated CRBN-AMPKα colocalization in IAV-infected cells Fig3C and S2B Fig.

**Figure 3.**
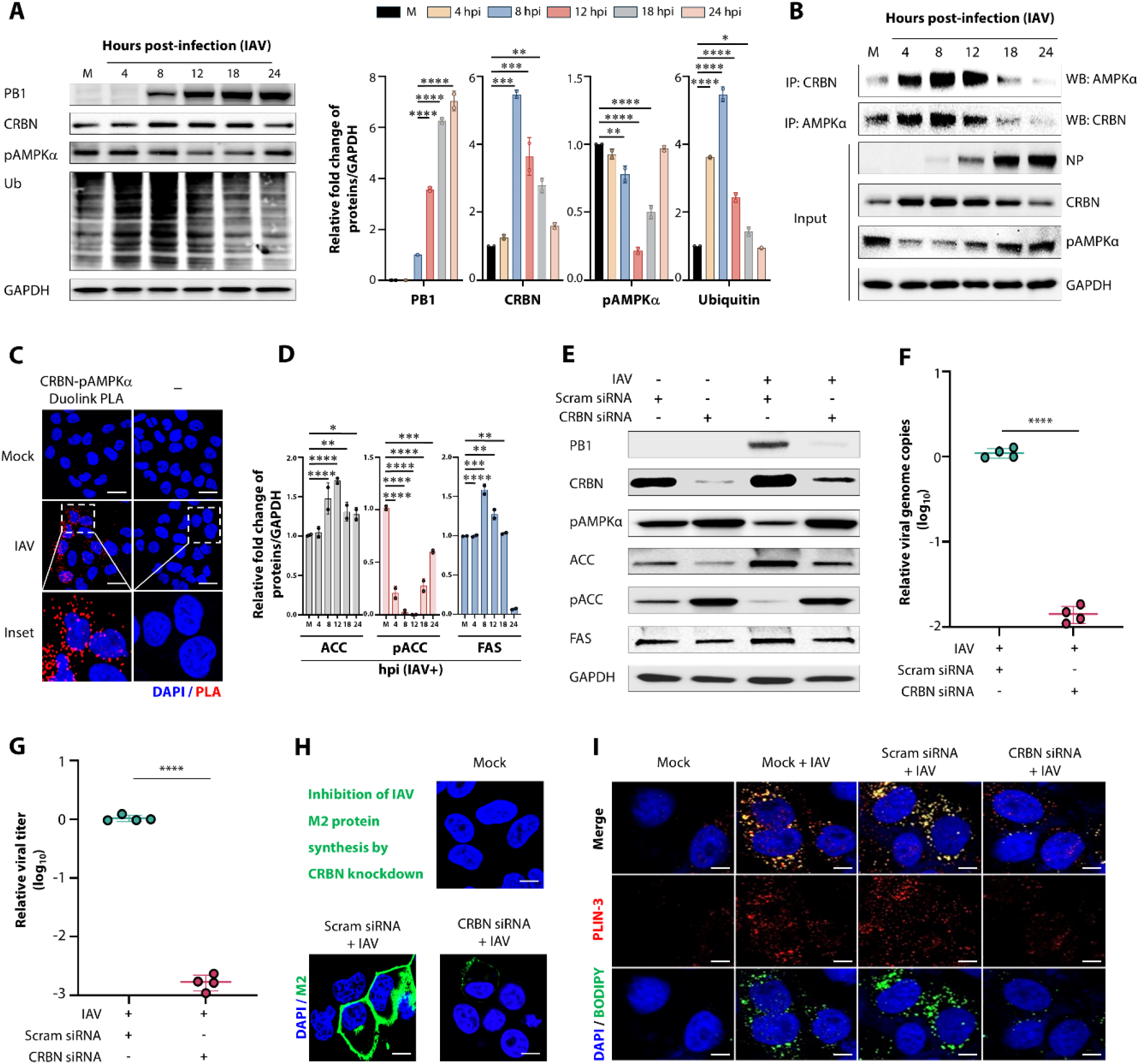
Suppression of IAV replication by blocking direct interaction between CRBN and AMPK. (A) Representative western blot images (left) and graphical representations (right) illustrating the sequential changes in total ubiquitinated proteins along with IAV PB1 protein, CRBN, and pAMPKα in the A549 cells infected with the IAV PR8 strain at an MOI of 1 FFU. GAPDH served as a loading control. (B) Representative immunoprecipitation assay results showing the direct interaction between CRBN and pAMPKα and vice versa in A549 cells infected with the IAV PR8 strain at an MOI of 1 FFU. GAPDH was used as a loading control. NP, IAV nucleoprotein. (C) Representative confocal images revealing the direct interaction between CRBN and pAMPKα in the A549 cells infected with the IAV PR8 strain at an MOI of 1 FFU by Duolink proximity assay. (D) Sequential changes of acetyl-CoA carboxylase (ACC), fatty acid synthase (FAS), and phosphorylated ACC (pACC) in the cells infected with IAV PR8 strain at an MOI of 1 FFU by western blot analysis. GAPDH served as a loading control. (E) Representative western blot images. Knockdown of CRBN reduced expression levels of IAV PB1 protein, CRBN, ACC, and FAS but increased pAMPKα and pACC. GAPDH served as a loading control. (F and G) Graphical representation of relative viral genome copies (F) and viral titers (G). Knockdown of CRBN by the transfection with siRNA against the *Crbn* gene in the A549 cells infected with the IAV PR8 strain at an MOI of 1 FFU reduced viral genome copy numbers and titers of IAV compared to the scrambled siRNA-transfected control. (H) Representative confocal images illustrating the inhibition of IAV M2 protein synthesis in response to the silencing of the *Crbn* gene in the A549 cells infected with the IAV PR8 strain at an MOI of 1 FFU. (I) Representative confocal images. Compared to scrambled siRNA-transfected IAV-infected control, silencing of CRBN decreased the number of lipid droplets (LDs) observed as colocalized spots of LD-associated perilipin 3 (PLIN-3, red) and BODIPY-stained LDs (green) in IAV-infected cells (G, PR8 strain, MOI of 1 FFU). Results are presented as arithmetic means ± S.D. Differences were evaluated using a one-way ANOVA. \**p* < 0.05; \*\**p* < 0.001; \*\*\**p* < 0.0001; \*\*\*\**p* < 0.00001. The scale bars in panels C, H, and I correspond to 20 µm, 10 µm, and 5 µm, respectively.

Since CRBN-induced AMPK degradation promotes anabolic metabolism, we next examined the effects of CRBN on fatty acid biosynthesis and LD formation during viral infection. The expression dynamics of acetyl-CoA carboxylase (ACC) and fatty acid synthase (FAS), both key enzymes in fatty acid biosynthesis, increased in parallel with CRBN levels but decreased as pAMPKα levels rose Fig 3D and S3 Fig. Similarly, phosphorylated ACC (pACC), an AMPK substrate that inhibits fatty acid biosynthesis [40] displayed an inverse expression pattern to CRBN and lipogenic enzymes Fig 3D and S3 Fig.

To further validate this regulator axis, we knocked down CRBN and observed increased pAMPKα and pACC levels, accompanied by reduced ACC and FAS expression Fig 3E. CRBN depletion also led to significant suppression of viral protein synthesis, genome replication, and progeny production Figure 3E-H, likely due to inhibition of LD formation in IAV-infected cells Fig 3I. Taken together, these findings suggest that CRBN promotes AMPK degradation early in infection, shifting cellular metabolism toward fatty acid biosynthesis and LD formation, ultimately facilitating viral replication at later stages [17, 34].

### *In Vitro* Broad-Spectrum Antiviral and Safety Profile of CRBN Blockers

To determine whether pharmacological inhibition of CRBN mimics the effects of CRBN knockdown on LD formation and viral replication, we tested thalidomide, a well-characterized immunomodulatory imide drug (IMiD) that exists as a racemic mixture of (*R*)-thalidomide and (*S*)-thalidomide [41, 42]. Treatment of IAV-infected cells with thalidomide (a mixture of two enantiomers *R* and *S*) or (*R*)-thalidomide reduced CRBN expression in a dose-dependent manner, leading to the activation of pAMPKα and pACC Fig 4A and S4A-B Fig. This, in turn, suppressed viral protein synthesis, viral genome replication, and progeny virus production Fig 4A-D and S4B-E Fig. Further analysis revealed that chemical inhibition of CRBN with thalidomide and its derivatives (lenalidomide or pomalidomide) or genetic silencing of CRBN significantly reduced the number of LDs in the cells infected with IAV, BCoV, PEDV, RVA, PRRSV, and PSaV in the mid-stage of infection Fig 4E and S5 Fig. Notably, inhibition of LD formation by thalidomide or (*R*)-thalidomide was accompanied by a reduction in IAV hemagglutinin (HA) palmitoylation Fig 4F and S4F Fig, a decrease in mitochondrial energy production at later stages of infection S4G Fig, and viral replication compartments of BCoV S4H Fig in the late stage of infection, further supporting the role of CRBN in virus-induced metabolic reprogramming [17, 43, 44].

**Figure 4.**
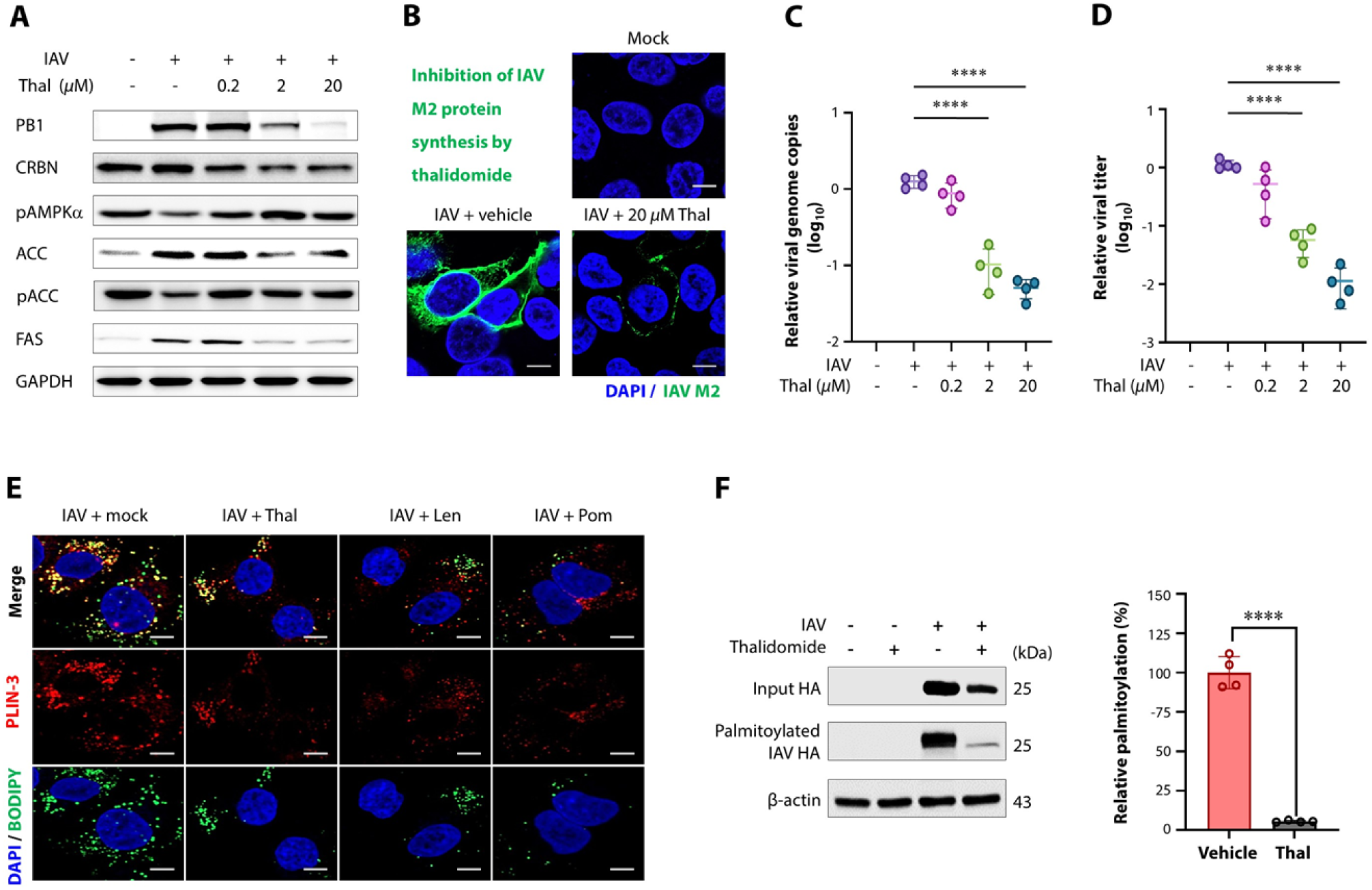
Chemical inhibition of CRBN suppresses IAV replication by preventing the LD formation. (A) Representative western blot images illustrating the suppression of IAV PB1 protein synthesis through inhibition of CRBN and associated lipogenic enzymes ACC and FAS, but activation of pAMPK and pACC by treatment of IAV-infected cells (PR8 strain, MOI = 1 FFU) with thalidomide in a dose-dependent manner. GAPDH served as a loading control. (B-D) Suppression of IAV protein synthesis (B), genome copy numbers (C), and progeny virus production (D) by treatment of IAV-infected cells (PR8 strain, MOI = 1 FFU) with 0.2 µM, 2 µM, and 20 µM of thalidomide. IAV M2, IAV matrix protein 2. (E) Representative confocal images. Compared to mock-treated IAV-infected control, inhibition of CRBN by the treatment of 20 µM of thalidomide, lenalidomide, and pomalidomide decreased the number of lipid droplets (LDs) as observed as colocalized spots of LD-associated perilipin 3 (PLIN-3, red) and BODIPY-stained LDs (green) in IAV-infected cells (E, PR8 strain, MOI of 1 FFU). (F) Representative western blot (left) and quantification (right) of inhibitory effects of thalidomide on palmitoylations of HA protein in IAV-infected cells (MOI = 1 FFU) at 24 hpi. Results are presented as arithmetic means ± S.D. Differences were evaluated using a one-way ANOVA. \**p* < 0.05; \*\**p* < 0.001; \*\*\**p* < 0.0001; \*\*\*\**p* < 0.00001. The scale bars in panels B and E correspond to 10 µm and 5 µm, respectively.

To assess the antiviral potency and cytotoxicity of (*R*)-thalidomide, we determined its half-maximal cytotoxicity concentration (CC_50_) and half-maximal inhibitory concentration (IC_50_) against various IAV and IBV strains, including pandemic strain S1 and S2 Tables. The CC_50_ in MDCK cells was 207.2 µM, while the IC_50_ values across different IAV and IBV strains remained in the low micromolar range S6 Fig, yielding selectivity index (SI) values ranging from 37 (for IBV Austria, Victoria lineage) to 103 (for IAV Korea H9N2) S6 Fig. Comparable antiviral effects were observed for BCoV, PEDV, RVA, PRRSV, and PSaV S7 Fig. These findings indicate that (*R*)-thalidomide exhibits broad-spectrum antiviral activity against seasonal IAV and IBV and pandemic IAV strains, reinforcing the potential of CRBN inhibition as a host-directed broad-spectrum antiviral strategy.

### Protection of IAV-Infected Mice by Immunomodulatory Imide Drugs (IMiDs)

Next, we assessed the *in vivo* antiviral efficacy of IMiDs, particularly (*R*)-thalidomide, in a mouse model infected with the mouse-adapted PR8 (H1N1) strain. The administration of (*R*)-thalidomide to IAV-challenged mice reduced mortality, body weight loss, and clinical scores in a dose-dependent manner, achieving 53.3% survival at a dose of 5 mg kg^-1^ d^-1^ Fig 5A-D. (*R*)-thalidomide significantly decreased IAV replication in the lungs, alleviating severe IAV-challenged lung lesions Fig 5E-G. Notably, (*R*)-thalidomide treatment suppressed IAV-induced CRBN expression, maintaining pAMPKα levels comparable to those in uninfected mice Fig 5E. The restoration of AMPK activity lowered lipogenic enzymes ACC and FAS to basal levels or below Fig 5E, suppressed viral RNA synthesis Fig 5F, and reduced progeny virus production Fig 5G, ultimately decreasing lung pathology and viral antigen distribution Fig 5H. These findings demonstrate that CRBN inhibition by IMiDs, such as (*R*)-thalidomide, protects mice from lethal IAV infection by sustaining AMPK activity and blocking virus-induced lipogenesis, a key process in viral replication.

**Figure 5.**
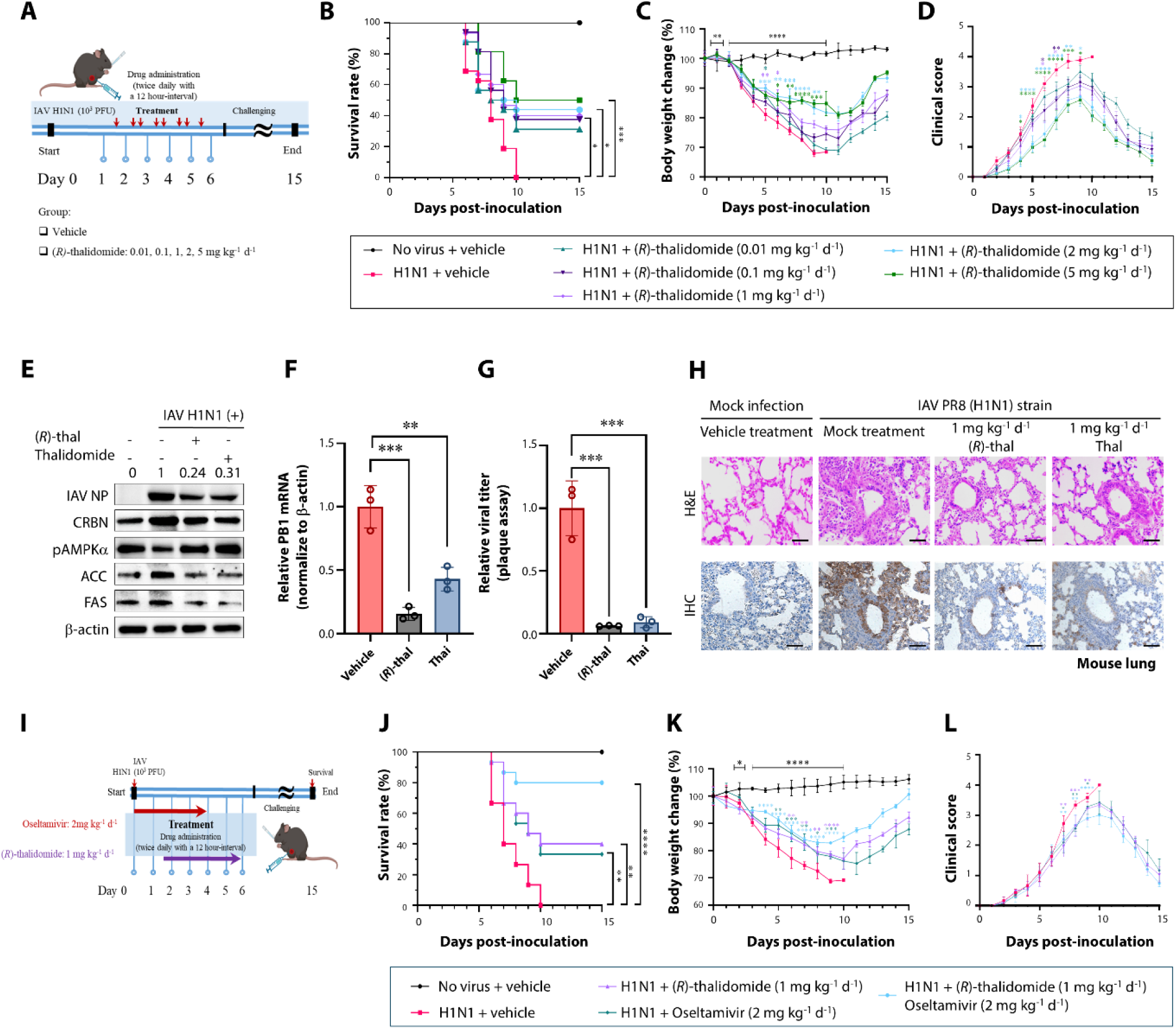
*In vivo* antiviral effects of (*R*)-thalidomide against influenza A virus (IAV) (A) Administration scheme of (*R*)-thalidomide; Mice (n = 16) were given various doses of (*R*)-thalidomide for four consecutive days, starting 1.5 days after being challenged with 10^3^ PFU of the mouse-adapted IAV PR8 (H1N1) strain. (B-D) Survival rates (expressed as percentages), daily body weight changes, and daily clinical scores of mice challenged with 10^3^ PFU of mouse-adapted IAV PR8 strain and treated with different doses of (*R*)-thalidomide. (E-G) Reduction in IAV protein synthesis (E), genome copy number (F), and infectious progeny production (G) in lungs sampled from IAV-challenged mice (n = 5) treated with 1 mg kg^-1^ d^-1^ (*R*)-thalidomide and thalidomide. (H) Representative images illustrating histological lesions (upper panels) and antigen distribution (lower panels) in the lung of mock- or IAV-infected mice, treated with either vehicle, 1 mg kg^-1^ d^-1^ (*R*)-thalidomide, or 1 mg kg^-1^ d^-1^ thalidomide. Amelioration of histological lung lesions (upper panels) and inhibition of IAV replication (lower panels) treated with (*R*)-thalidomide or thalidomide. (I) Administration scheme of combination therapy: Mice (n = 16) were administered the 1 mg kg^-1^ d^-1^ (*R*)-thalidomide or 2 mg kg^-1^ d^-1^ oseltamivir, either individually or in combination, twice daily for four consecutive days, 1.5 days after challenge with 10^3^ PFU of mouse-adapted IAV PR8 (H1N1) strain. (J-L) Survival rates (expressed as percentages), daily body weight changes, and daily clinical scores of mice challenged with 10^3^ PFU of mouse-adapted IAV PR8 strain and treated by combination therapy. Results are presented as arithmetic means ± S.D. \**P* < 0.05; \*\**P* < 0.01; \*\*\**P* < 0.001, \*\*\*\**p* < 0.00001, a one-way analysis of variance with Tukey’s correction for multiple comparisons. The scale bar of panel H corresponds to 800 µm.

### Enhanced Antiviral Effect of Combination Therapy with Direct-Acting Oseltamivir

A combination therapy of antiviral drugs with different mechanisms of action can increase antiviral efficacy [45–49]. We examined the antiviral efficacy of thalidomide and oseltamivir in various ratios to find the combination therapy with the best effect Fig 5I-L and S8 Fig. Treatment with 2 mg kg^-1^ d^-1^ oseltamivir combined with 1 mg kg^-1^ d^-1^ (*R*)-thalidomide resulted in an 80.0% survival rate with lesser body weight loss and clinical scores Fig 5I-L and S8 Fig, compared to 40.0% survival with 1 mg kg^-1^ d^-1^ (*R*)-thalidomide alone and 33.3% survival with 2 mg kg^-1^ d^-1^ oseltamivir alone Fig 5I-L and S8 Fig. These findings indicate that combining low doses of antiviral agents with complementary mechanisms of action may enhance antiviral efficacy and improve outcomes.

## Discussion

During viral infection, host cells often shift their intracellular microenvironment from a catabolic to an anabolic state, enhancing LD formation to support viral replication [17, 22, 23]. This lipidomic reprogramming suggests an early inhibition of AMPK, a key regulator of catabolism [27, 29, 32, 33]. Our study discovered a dynamic suppression of pAMPKα from the early to mid-stage infection, which later recovers across multiple RNA viruses. Notably, CRBN expression exhibited an inverse correlation with AMPKα levels, strongly suggesting a regulatory interaction. Immunoprecipitation and Duolink PLA assays confirmed that CRBN directly binds to AMPKα early in infection, likely facilitating AMPKγ ubiquitination and degradation, thereby suppressing AMPK activity via the CRBN-mediated ubiquitin-proteasome pathway [26–31].

CRBN-driven AMPK degradation in virus-infected cells appears to establish an anabolic microenvironment, coinciding with SREBP-dependent lipidomic reprogramming [22, 50]. Our data show that ACC and FAS, key lipogenic enzymes regulated by SREBP-1, mirror CRBN expression dynamics while being inversely correlated with those of pAMPKα [27, 29, 32, 33]. CRBN inhibition, either via siRNA or chemical blockade, prevented LD formation and restored AMPK activation, shifting cellular metabolism toward a catabolic state. Given that AMPK directly phosphorylates SREBP-1c and SREBP-2 [51, 52] inhibiting their nuclear translocation and downstream lipogenic gene transcription [51, 52] our findings suggest that early AMPK degradation by CRBN facilitates SREBP-driven lipogenesis and LD formation, crucial for viral replication [22, 50–52].

Thalidomide has been reported to neither affect AMPK activation nor alter CRBN-AMPKα binding affinity, suggesting that CRBN regulates AMPK through a pathway independent of the CRBN-IMiD binding region [53]. However, our findings contradict this hypothesis, as thalidomide treatment restored pAMPKα levels in virus-infected cells. Moreover, thalidomide and its derivatives significantly reduced LD formation, supporting the idea that IMiDs inhibit CRBN, thereby activating AMPK [30, 54, 55]. Further studies are needed to dissect the precise molecular mechanisms by which thalidomide influences CRBN-mediated AMPK regulation in the context of viral infection.

Many RNA viruses exploit host lipid metabolism at multiple stages of their life cycle to enhance entry, replication, and egress [25, 35–39]. Early LD accumulation often compensates for cellular nutritional deficiencies due to cell membrane damage at later stages [17–21]. Our data confirmed that RNA viruses, including IAV, IBV, PEDV, BCoV, RVA, PRRSV, and PSaV, utilize a conserved strategy to degrade AMPK complexes via CRBN, shifting host metabolism from catabolism to anabolism [14–16]. The concurrent activation of ACC and FAS with CRBN expression suggests that CRBN-mediated AMPK inhibition serves as a key trigger for SREBP-dependent metabolic reprogramming [22, 50]. These findings highlight CRBN as a conserved metabolic switch hijacked by diverse RNA viruses to promote lipid synthesis and LD formation, thereby optimizing replication conditions [17, 22–25]. Importantly, these findings ultimately led to the discovery that IMiDs, which are CRBN inhibitors, could be used as host-directed broad-spectrum antiviral therapeutics for various RNA virus infections [10].

Thalidomide and its derivatives have been used to treat various diseases, including multiple myeloma, erythema nodosum leprosum, autoimmune conditions, and solid tumors [56–59]. However, due to the severe birth defects and impact on fetal development, they cannot be used by pregnant women and women of childbearing age, therefore limiting their widespread use [60]. Combination therapy provides a potential solution to reduce the required dosage and teratogenic risks of these compounds while maintaining their antiviral activity [45–49]. In this study, a combination therapy of a low dose of thalidomide with the NA inhibitor, oseltamivir, showed a more potent antiviral effect than each treatment. However, our work rather highlights the critical role of the CRBN-AMPK-LD pathways in the life cycles of multiple viruses, suggesting the potential to identify broad-spectrum approaches.

In summary, our study identifies CRBN as a critical regulator of RNA virus replication by degrading AMPK and promoting metabolic reprogramming Fig 6. By shifting the intracellular environment from catabolism to anabolism, CRBN promotes LD accumulation, providing essential substrates for viral replication. While IMiDs effectively inhibit CRBN and suppress viral replication, their well-documented teratogenic effects limit their clinical utility as antiviral therapeutics. However, our findings offer key insights into an essential virus-host interaction that could be harnessed for therapeutic intervention. Future efforts should focus on identifying novel, non-teratogenic molecules targeting the CRBN-AMPK axis, offering a safer, more feasible strategy for host-directed antiviral therapy against influenza and other RNA viral infections.

**Figure 6.**
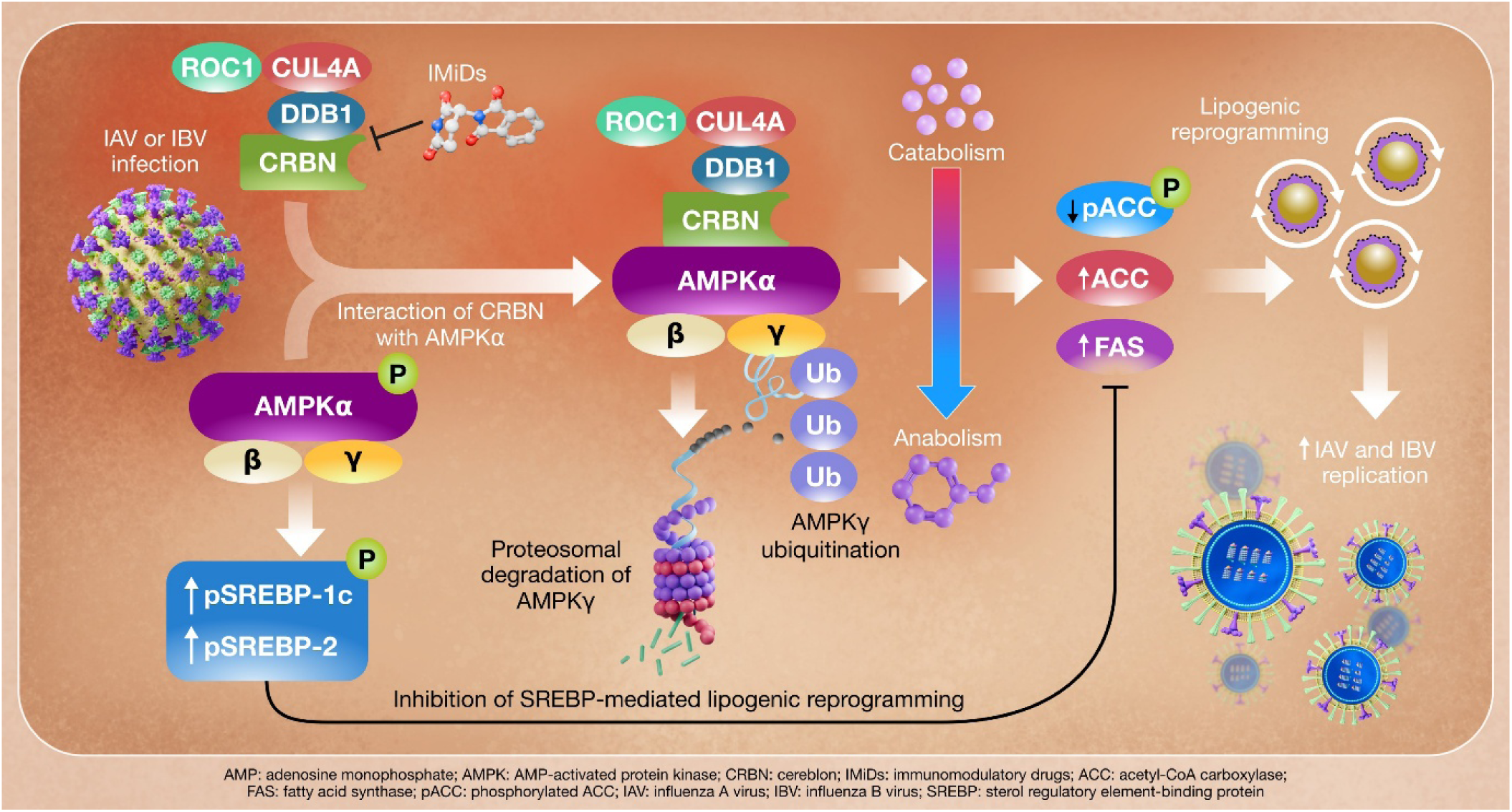
Schematic diagram. Cereblon (CRBN) facilitates the replication of types A and B influenza viruses (IAV and IBV) by ubiquitinating AMPK. Infection with IAV or IBV leads to binding of CRBN and AMPKα, resulting in the ubiquitination and proteasomal degradation of AMPKγ. As a result, the cell environment shifts from catabolism to anabolism, activating intracellular lipogenic enzymes and inducing intracellular lipogenic reprogramming. In the normal microenvironment, AMPK complexes phosphorylate SREBP-1c and SREBP-2, which prevents SREBP-mediated lipogenic reprogramming. However, when the CRBN-ubiquitin proteasome system degrades AMPKγ, lipogenic enzymes such as ACC and FAS are induced, leading to intracellular lipogenic reprogramming. Ultimately, intracellular lipogenic reprogramming, prompted by virus-induced early activation of the CRBN-AMPK axis, facilitates the replication of IAV and IBV. Inhibition of CRBN in virus-infected cells by treating with immunomodulatory drugs (IMiDs) prevents lipid droplet formation and restores AMPK activation, shifting cellular metabolism toward a catabolic state. Therefore, CRBN inhibitors could act as host-directed pan-influenza antivirals.

## Materials and methods

### Animal ethics statement

All procedures were conducted in accordance with the requirements set forth by the Institutional Animal Care and Use Committees at Chonnam National University (CNU IACUC-YB-2018-48 and CNU IACUC-YB-2023-171,176). The care and handling of the animals adhered to all current international laws and regulations, including the NIH Guide for the Care and Use of Laboratory Animals (NIH Publication No. 85-23, 1985, revised 1996). All experiments were designed to minimize both the number of animals used and their suffering.

### Cells and viruses

Vero E6, MDCK, LLC-PK, MA104, A549, Caco-2, MARC-145, and HRT-18G cells were cultured in EMEM, α*-*MEM, or DMEM *at* 37°C in 5% CO_2_. All media were supplemented with 10% fetal bovine serum, 100 U/mL penicillin, and 100 μg/mL streptomycin. The strains of IAV used included Puerto Rico/8 (PR8) (H1N1), California/04/2009 (H1N1) (2009 Pandemic H1N1-like), Brisbane/10/07 (H3N2), and Chicken/Korea/01310/2001 (H9N2). The two strains of IBV used included Austria/1359417/2021 (B/Victoria) and Phuket/3073/2013 (B/Yamagata). Other viruses used, as stated in the text, were BCoV KWD20, PEDV QIAP1401, PRRSV LYM, RVA NCDV, and PSaV Cowden strains. Detailed procedures for cell and virus culture, as well as virus titration, are provided in the Supplementary Information.

### Reagents, kits, siRNAs, and antibodies

The details of the reagents, antibodies, siRNAs, and kits used in this study are provided in the Supplementary Information.

### Cytotoxicity assessment

The MTT assay was used to determine the half-maximal cytotoxic concentration (CC_50_) of the chemicals and their solvents, as described in the Supplementary Information.

### Duolink proximity ligation assay (DPLA)

The direct interaction of CRBN with AMPKα was determined using the Duolink PLA kit (Sigma-Aldrich), following established procedures [61].

### Transfection of siRNA

siRNAs targeting the *Crbn* gene (10 nmol) were transfected into A549 cells at 70% and 80% confluency, which were grown in 12-well culture plates or 8-well chamber slides, using Lipofectamine 3000 (Thermo Scientific) as previously described [61]. To optimize efficient knockdown of target proteins, a second transfection was performed 24 h after the first, and the cells were incubated for an additional 24 h. As a negative control, scrambled siRNA (10 nmol) was transfected as described above.

### Determination of fatty acid oxidation (FAO)

The colorimetric FAO assay kit (AssayGenie) was used to measure FAO activity in vehicle- or IMiD-treated cells, as described in the Supplementary Information.

### Treatment of inhibitory chemicals to cells

Cell lines were cultured in 6- or 12-well plates or 8-well chamber slides and washed twice with phosphate-buffered saline (PBS, pH 7.4) before infection at the following multiplicities of infection (MOI): MOI of 1 or 0.1 FFU/cell of IAV, BCoV, PEDV, PRRSV, RVA, and PSaV. The cells were allowed to absorb the virus and washed twice with PBS (pH 7.4). Different concentrations of IMiDs (0.2 μM, 2 μM, or 20 μM) or vehicle were added to cells immediately after virus adsorption and incubated for the indicated time. Cell lysates and supernatants were collected as described below and in the Supplementary Information section.

### Immunoprecipitation assay

CRBN and AMPKα co-immunoprecipitation was performed as described previously [32,61]. Briefly, A549 cells grown in 6-well plates were mock- or infected with the IAV PR8 strain at an MOI of 1 FFU and incubated for the indicated times at 37 °C. Cell lysates were pre*-*cleared by incubation with protein A- or G-agarose beads for 30 min at 4 °C and incubated with antibodies against CRBN or AMPKα overnight at 4 °C. Antibody–protein complexes were precipitated with equilibrated Protein A- or G-agarose beads at 4 °C for 3 h, incubated with lysis buffer at 37 °C for 15 min, and evaluated by western blot analysis, as described below.

### Experimental animals

CRBN KO mice were generated from C57BL/6N mice as described previously [33]. The genotype of CRBN WT and KO mice was determined by PCR and western blot analysis as described in the Supplementary Information. Seven-week-old female wild-type (WT) C57BL/6J mice were purchased from Samtako (Osan, South Korea) and used to assess survival following IAV infection, observe gross lung lesions, analyze viral replication, evaluate changes in host genes, and identify pathological changes in response to IAV infection and chemical treatments. The mice were housed in standard cages and exposed to a 12:12-hour light/dark cycle at 25°C, with food and water available *ad libitum* in a specific-pathogen-free facility. After 1 week of acclimatization, mice were used for each experiment as described below.

### In vivo organ-specific toxicity

Evaluation of IMiD organ-specific toxicity in animals was performed according to the procedures described in the Supplementary Information.

### Determination of median lethal dose (LD_50_) of mouse-adapted PR8 strain

The LD_50_ of the mouse-adapted PR8 strain were determined by infecting 8-week-old mice [17,62]. Detailed procedures and evaluation of IAV-induced clinical scores are described in Supplementary Information.

### In vivo antiviral activity

The antiviral effects of IMiDs, either alone or in combination with oseltamivir, were tested against IAV infection in a mouse model, as detailed in the Supplementsary Information.

### Median Tissue culture infectious dose (TCID_50_) assay

The TCID_50_ assay was conducted to measure the titer of PSaV, as detailed in the Supplementary Information.

### Immunofluorescence assay (IFA)

IFA was used to determine the dynamics of CRBN, BODIPY-stained LD, PLIN-1 and −3, viral antigens, and infectivity in cultured cells or lung tissue, as described in Supplementary Information.

### Western blot analysis

Western blot analysis was conducted to detect specific cellular or viral proteins in cultured cells or lung tissues. Normalization and graphic representation followed the protocols outlined in the Supplementary Information.

### Quantitative real-time PCR

Real-time quantitative PCR was performed to detect and quantify target viral RNAs and host mRNAs from cultured cells or animal samples, as outlined in the Supplementary Information.

### Palmitoylation assay

The impact of IMiDs on the palmitoylation of IAV HA in virus-infected cells was assessed using the CAPTUREome™ S-palmitoylated protein kit (Badrilla), as detailed in the Supplementary Information.

### Histopathology

As described in the Supplementary Information, histopathological changes were analyzed in WT or *Crbn* KO mice infected with or without IAV and either treated with IMiDs or vehicle.

### Immunohistochemistry (IHC)

The expression levels of IAV proteins in experimental animals were evaluated using IHC, as detailed in the Supplementary Information.

### Statistical analyses and software

Statistical analyses were performed on triplicate experiments using one-way ANOVA in GraphPad Prism version 10.3.1 (GraphPad Software Inc., La Jolla, CA, USA). *P* values of less than 0.05 were considered statistically significant. Figures were generated using Adobe Photoshop CS6 and Prism 10 version 10.3.1. *Illustrations:* Illustrations of mice were created using BioRender software (https://biorender.com/).

## Supporting information

**Table S1.** The median cytotoxic concentration (CC_50_) of (*R*)-thalidomide and thalidomide in different cell lines determined by measuring the cellular NAD(P)H-dependent cellular oxidoreductase enzymes (MTT assay)

| Cell line | CC<sub>50</sub> |  |
| --- | --- | --- |
|  | (*R*)-thalidomide | Thalidomide |
| A549 | 207.2 ± 2.46 | 291.6 ± 2.34 |
| MDCK | 220.5 ± 2.34 | 217.6 ± 2.47 |
| HRT-18G | 285.6 ± 2.45 | 216.1 ± 2.34 |
| Vero E6 | 156.1 ± 2.19 | 233.0 ± 2.37 |
| MA104 | 222.0 ± 2.35 | 281.9 ± 2.45 |
| MARC-145 | 244.5 ± 2.39 | 276.8 ± 2.44 |
| LLC-PK | 325.8 ± 2.51 | 226.7 ± 2.36 |

**Table S2.** The half maximal (50%) inhibitory concentration (IC_50_) and selectivity index (SI) of (*R*)-thalidomide and thalidomide for RNA viruses.

| Virus<sup>a</sup> | Strain | *(R)*-thalidomide |  | Thalidomide |  |
| --- | --- | --- | --- | --- | --- |
|  |  | IC<sub>50</sub> (μM) | SI (CC<sub>50</sub>/IC<sub>50</sub>) | IC<sub>50</sub> (μM) | SI (CC<sub>50</sub>/IC<sub>50</sub>) |
| IAV | California/04/2009 (H1N1) | 2.44 ± 0.39 | 85 | 2.61 ± 0.42 | 83 |
| IAV | Puerto Rico/8/1934 (H1N1) | 3.00 ± 0.48 | 69 | 2.91 ± 0.47 | 75 |
| IAV | Brisbane10/2007 (H3N2) | 3.69 ± 0.57 | 56 | 3.81 ± 0.58 | 57 |
| IAV | Chicken/Korea/01310/21001 (H9N2) | 2.01 ± 0.32 | 103 | 1.79 ± 0.25 | 122 |
| IBV | Phuket/3073/2013 (Yamagata) | 2.70 ± 0.43 | 77 | 2.58 ± 0.41 | 84 |
| IBV | Austria/1359417/2021 | 5.53 ± 0.74 | 37 | 4.17 ± 0.62 | 52 |
| BCoV | KWD20 | 1.20 ± 0.08 | 237 | 1.34 ± 0.13 | 161 |
| PEDV | QIAP1401 | 2.32 ± 0.37 | 67 | 2.50 ± 0.40 | 93 |
| RVA | NCDV | 2.21 ± 0.35 | 100 | 2.18 ± 0.34 | 129 |
| PRRSV | LMY | 2.45 ± 0.39 | 100 | 2.18 ± 0.34 | 127 |
| PSaV | Cowden | 27.27 ± 1.44 | 12 | 22.81 ± 1.36 | 10 |

<sup>a</sup>IAV, influenza A virus; IBV, influenza B virus; BCoV, bovine coronavirus; PEDV, porcine epidemic diarrhea coronavirus; RVA, species A
rotavirus; PRRSV, porcine reproductive and respiratory syndrome virus.

**S3 Table. Cell culture information.**

**S4 Table. Virus information.**

**S5 Table. Virus culture information.**

**S6 Table. Chemicals, kits, and siRNAs used in this study.**

**S7 Table. Antibodies used in this study.**

**S8 Table. Experimental design for determining organ-specific toxicity of (*R*)-thalidomide in the mice.**

**S9 Table. Experimental design for determining antiviral effects of (*R*)-thalidomide on influenza A virus infection in the mouse model.**

**S10 Table. Experimental design for determining combination therapy of (*R*)-thalidomide and oseltamivir on influenza A virus infection in the mouse model.**

**S11 Table. Experimental design for determining inhibitory effects of (*R*)-thalidomide on virus replication, and histopathological lesions in the lungs from IAV-infected mice.**

**S12 Table. Primers used for the detection of viral and host genes.**

**Figure S1.**
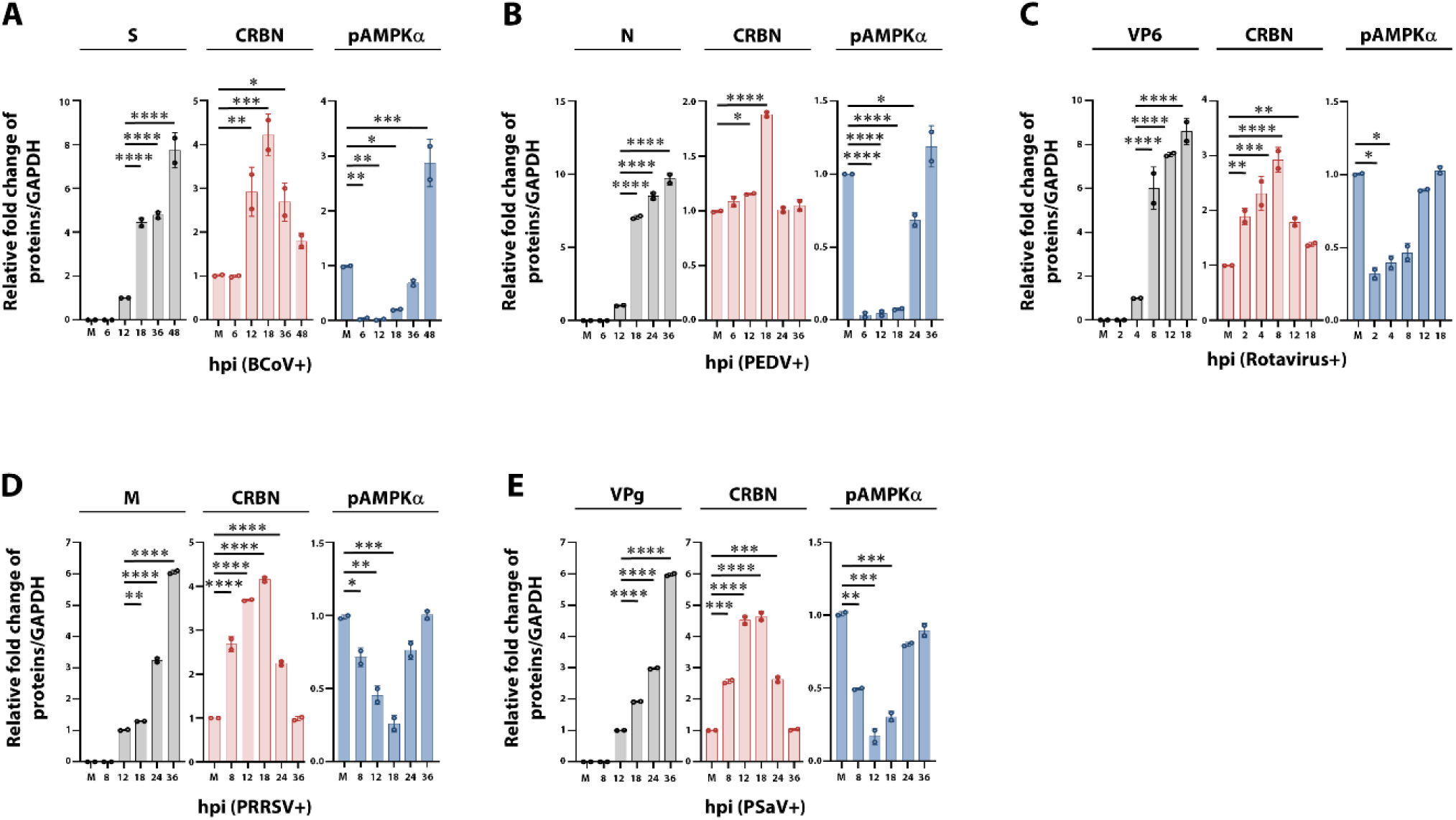
The dynamic pattern of CRBN and pAMPK*α* is common in cells infected with diverse RNA viruses. (A-E) Graphical representation of sequential changes of CRBN, pAMPKα, and each viral protein in the HRT-18G cells infected with bovine coronavirus (BCoV) KWD strain at an MOI of 0.1 FFU (A), Vero E6 cells infected with porcine epidemic diarrhea coronavirus (PEDV) QIAP1401 strain at an MOI of 0.1 FFU (B), MA104 cells infected with bovine species A rotavirus (RVA) NCDV strain at an MOI of 0.1 FFU (C), MARC-145 cells infected with porcine reproductive and respiratory syndrome virus (PRRSV) LMY strain at an MOI of 0.1 FFU (D), and LLC-PK cells infected with porcine sapovirus (PSaV) Cowden strain at an MOI of 0.1 FFU (E). S, BCoV spike protein; N, PEDV nucleocapsid protein; VP6, RVA capsid middle layer protein; M, PRRSV membrane protein; and VPg, PSaV genome-linked viral protein. Results are presented as arithmetic means ± S.D. Differences were evaluated using a one-way ANOVA. \**p* < 0.05; \*\**p* < 0.001; \*\*\**p* < 0.0001; \*\*\*\**p* < 0.00001.

**Figure S2.**
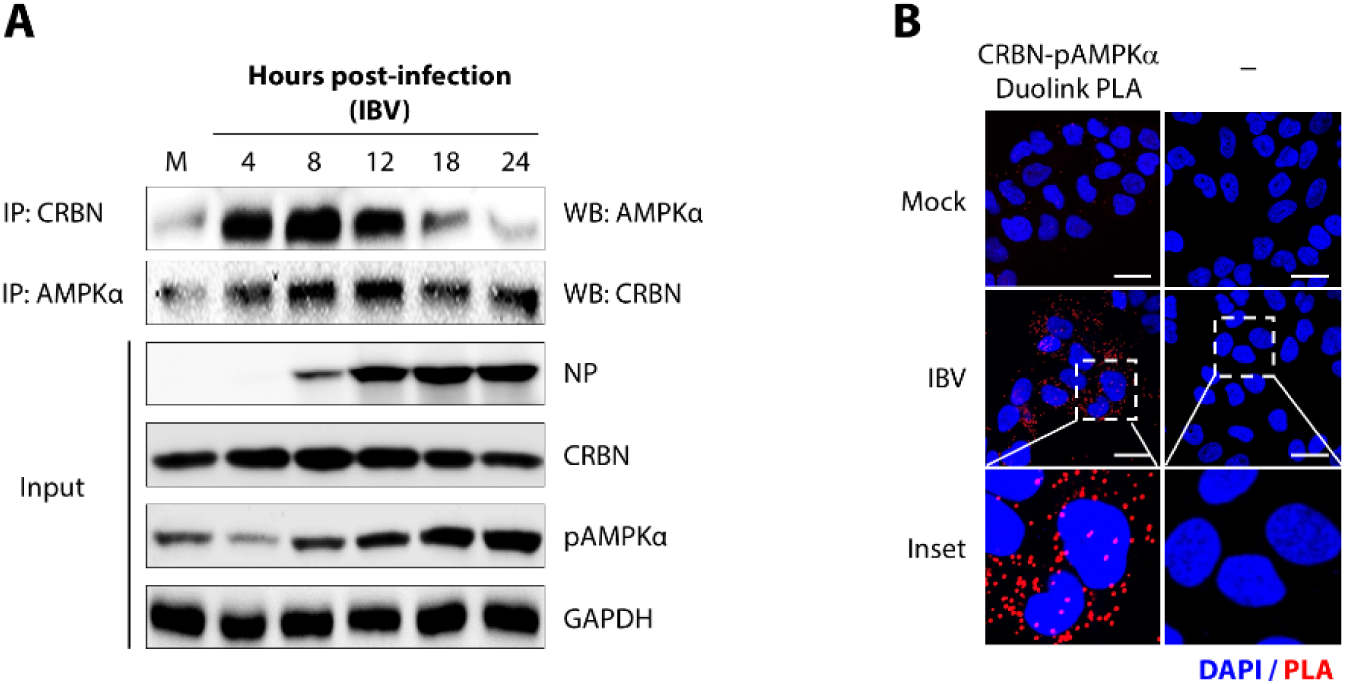
Direct interaction between CRBN and AMPK in the cells infected with the influenza B virus (IBV) (A) Representative immunoprecipitation assay results showing the direct interaction between CRBN and pAMPKα and vice versa in the A549 cells infected with the IBV Phuket strain at an MOI of 1 FFU. GAPDH was used as a loading control. NP, IBV nucleoprotein. (B) Representative confocal images revealing the direct interaction between CRBN and pAMPKα in the A549 cells infected with the IBV Phuket strain at an MOI of 1 FFU by Duolink proximity assay. The scale bars in panel B correspond to 20 µm.

**Figure S3.**
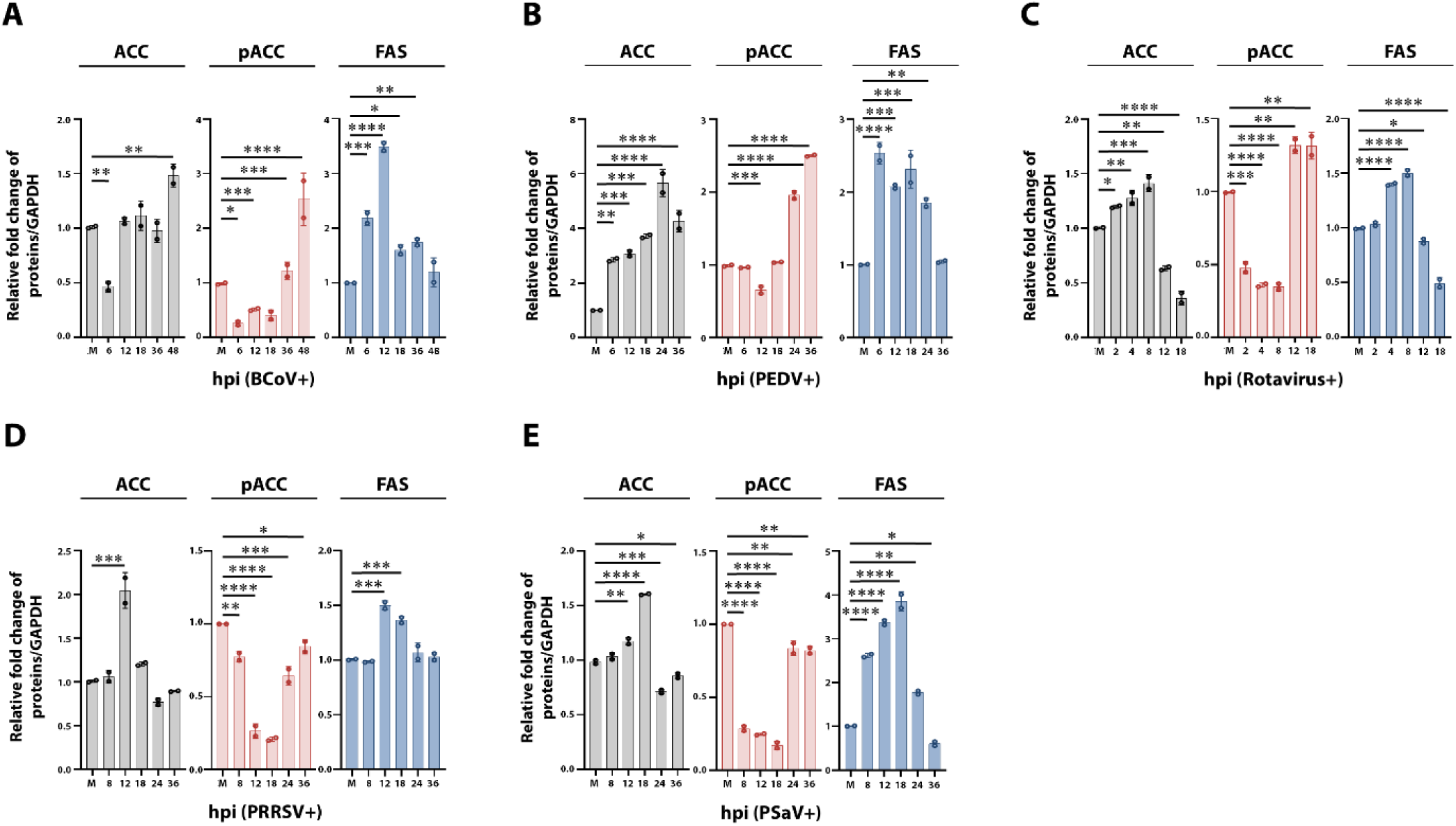
Upregulation of lipogenic enzymes due to the CRBN-associated degradation of AMPK complexes during RNA virus infections. (A-E) Graphical representation of sequential changes of lipogenic enzymes acetyl-CoA carboxylase (ACC) and fatty acid synthase (FAS), and its inhibitor phosphorylated ACC (pACC) in the cells infected with bovine coronavirus (BCoV) KWD strain at an MOI of 0.1 FFU (A), Vero E6 cells infected with porcine epidemic diarrhea coronavirus (PEDV) QIAP1401 strain at an MOI of 0.1 FFU (B), MA104 cells infected with bovine species A rotavirus (RVA) NCDV strain at an MOI of 0.1 FFU (C), MARC-145 cells infected with porcine reproductive and respiratory syndrome virus (PRRSV) LMY strain at an MOI of 0.1 FFU (D), and LLC-PK cells infected with porcine sapovirus (PSaV) Cowden strain at an MOI of 0.1 FFU (E). S, BCoV spike protein; N, PEDV nucleocapsid protein; VP6, RVA capsid middle layer protein; M, PRRSV membrane protein; and VPg, PSaV genome-linked viral protein. Results are presented as arithmetic means ± S.D. Differences were evaluated using a one-way ANOVA. \**p* < 0.05; \*\**p* < 0.001; \*\*\**p* < 0.0001; \*\*\*\**p* < 0.00001.

**Figure S4.**
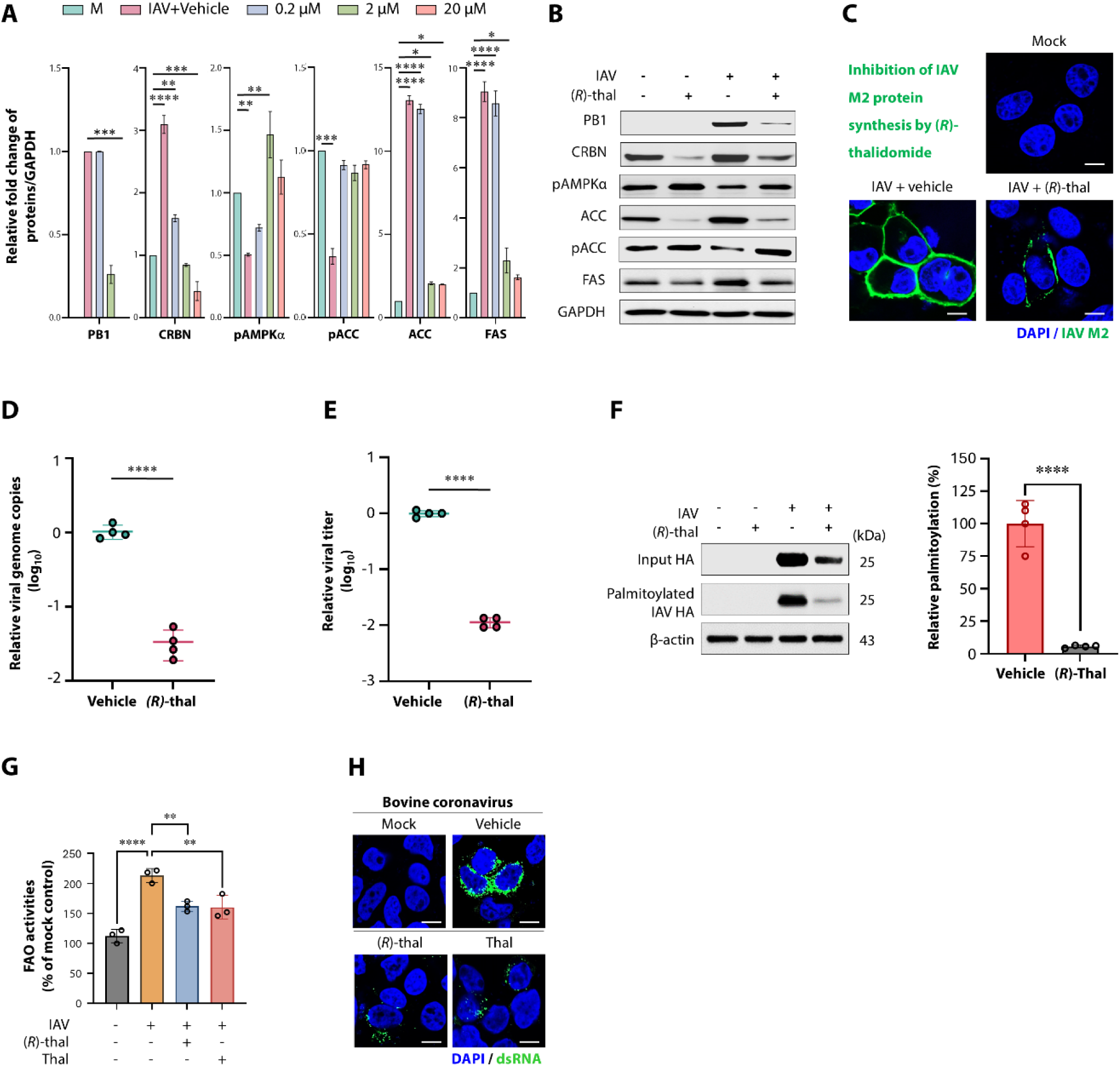
Inhibition of CRBN by treatment with thalidomide or (*R*)-thalidomide suppresses IAV replication. (A) Graphical representation showing reduction in IAV PB1 protein, CRBN, ACC, and FAS but activation of pAMPKα and pACC in the A549 cells infected with IAV PR8 strain at an MOI of 1 FFU by treatment with thalidomide in a dose-dependent manner. (B) Representative western blot images illustrating the suppression of IAV PB1 protein synthesis through inhibition of CRBN and associated lipogenic enzymes ACC and FAS, but activation of pAMPK and pACC by treatment of IAV-infected cells (PR8 strain, MOI = 1 FFU) with (*R*)-thalidomide in a dose-dependent manner. GAPDH served as a loading control. (C) Representative confocal images revealing a reduction in IAV matrix 2 (M2) protein in the A549 cells infected with the PR8 strain at an MOI of 1 FFU by treatment with 20 µM of (*R*)-thalidomide. (D-E) Suppression of viral genome replication (E) and progeny virus production (E) by treatment of IAV-infected cells (PR8 strain, MOI = 1 FFU) with 20 µM of (*R*)-thalidomide. (F) Representative western blot (left) and quantification (right) of inhibitory effects of (*R*)-thalidomide on palmitoylations of HA protein in IAV-infected cells (MOI = 1 FFU) at 24 hpi. (G) Graphical representation of inhibitory effects of (*R*)-thalidomide and thalidomide (mixture of two enantiomers, (*R*)-thalidomide and (*S*)-thalidomide) on intracellular fatty acid oxidation (FAO) activities in the IAV-infected cells (MOI = 1 FFU) at 24 hpi. (H) Inhibition of double-stranded RNA (dsRNA) formation in the bovine coronavirus-infected cells by the treatment with (*R*)-thalidomide and thalidomide. Note the relatively small amount of dsRNA-positive fluorescence signals in the (*R*)-thalidomide- and thalidomide-treated cells compared to vehicle-treated control cells. Results are presented as arithmetic means ± S.D. Differences were evaluated using a one-way ANOVA. \**p* < 0.05; \*\**p* < 0.001; \*\*\**p* < 0.0001; \*\*\*\**p* < 0.00001. The scale bars in panels C and H correspond to 10 µm and 13 µm, respectively.

**Figure S5.**
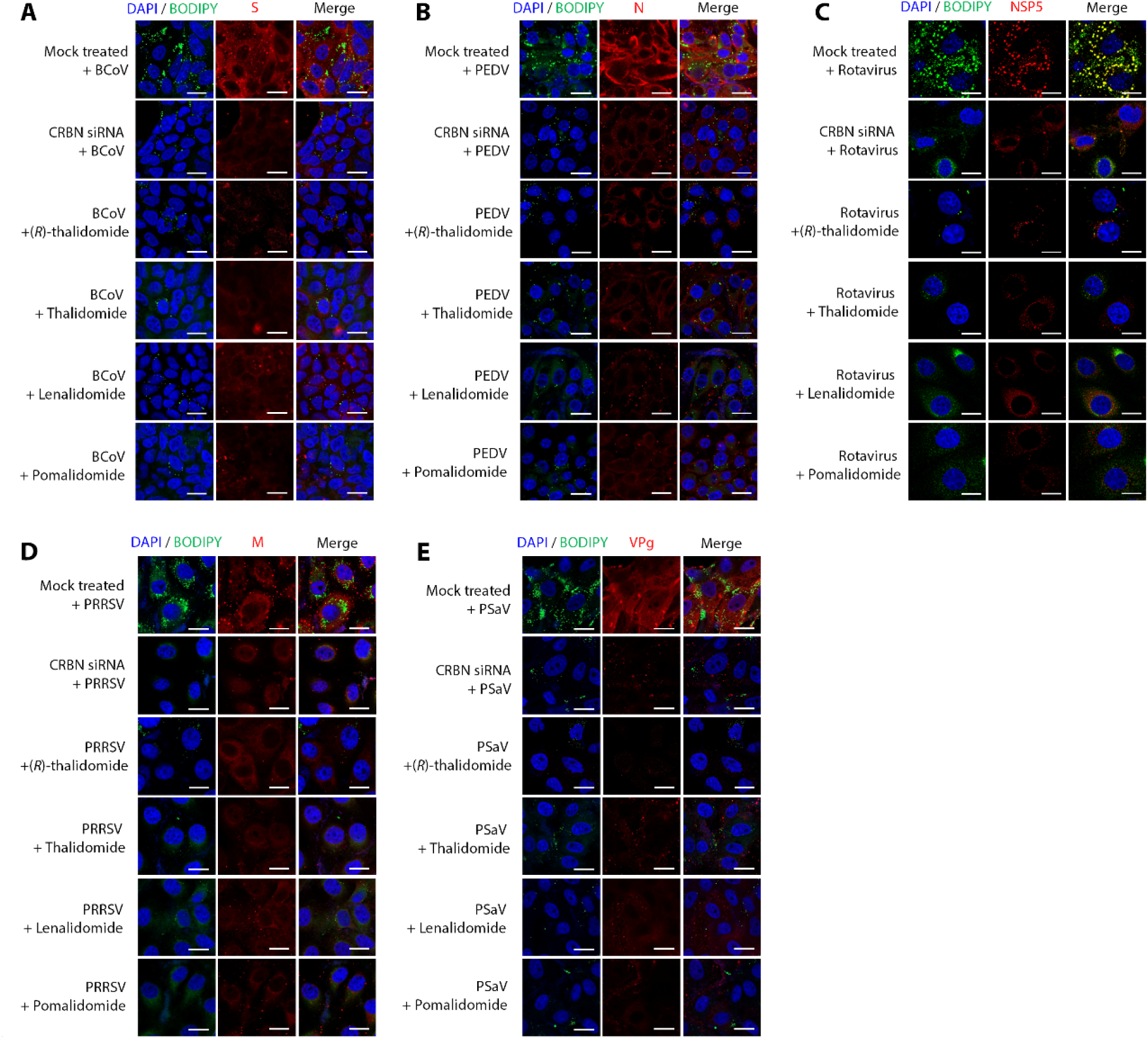
Inhibition of CRBN by IMiD treatment or Crbn siRNA transfection suppresses both LD formation and viral replication. (A-E) Representative confocal images. Compared to mock-treated virus-infected control, inhibition of CRBN by the treatment of 20 µM of (*R*)-thalidomide, thalidomide, lenalidomide, and pomalidomide or transfection with siRNA against *Crbn* gene decreased the number of BODIPY-stained LDs (green) and suppressed the expression levels of BCoV spike (S) protein (red) in BCoV-infected HRT-18G cells (A, KWD strain, MOI of 0.1 FFU), PEDV nucleocapsid (N) protein (red) in PEDV-infected Vero E6 cells (B, QIAP1401 strain, MOI of 0.1 FFU), RVA nonstructural protein 5 (NSP5) protein (red) in RVA-infected MA-104 cells (C, NCDV strain, MOI of 0.1 FFU), PRRSV membrane (M) protein (red) in PRRSV-infected MA-104 cells (D, LMY strain, MOI of 0.1 FFU), and PSaV viral-linked protein (VPg) protein (red) in PSaV-infected LLC-PK cells (E, Cowden strain, MOI of 0.1 FFU). The scale bars in panels A-E correspond to 10 µm.

**Figure S6.**
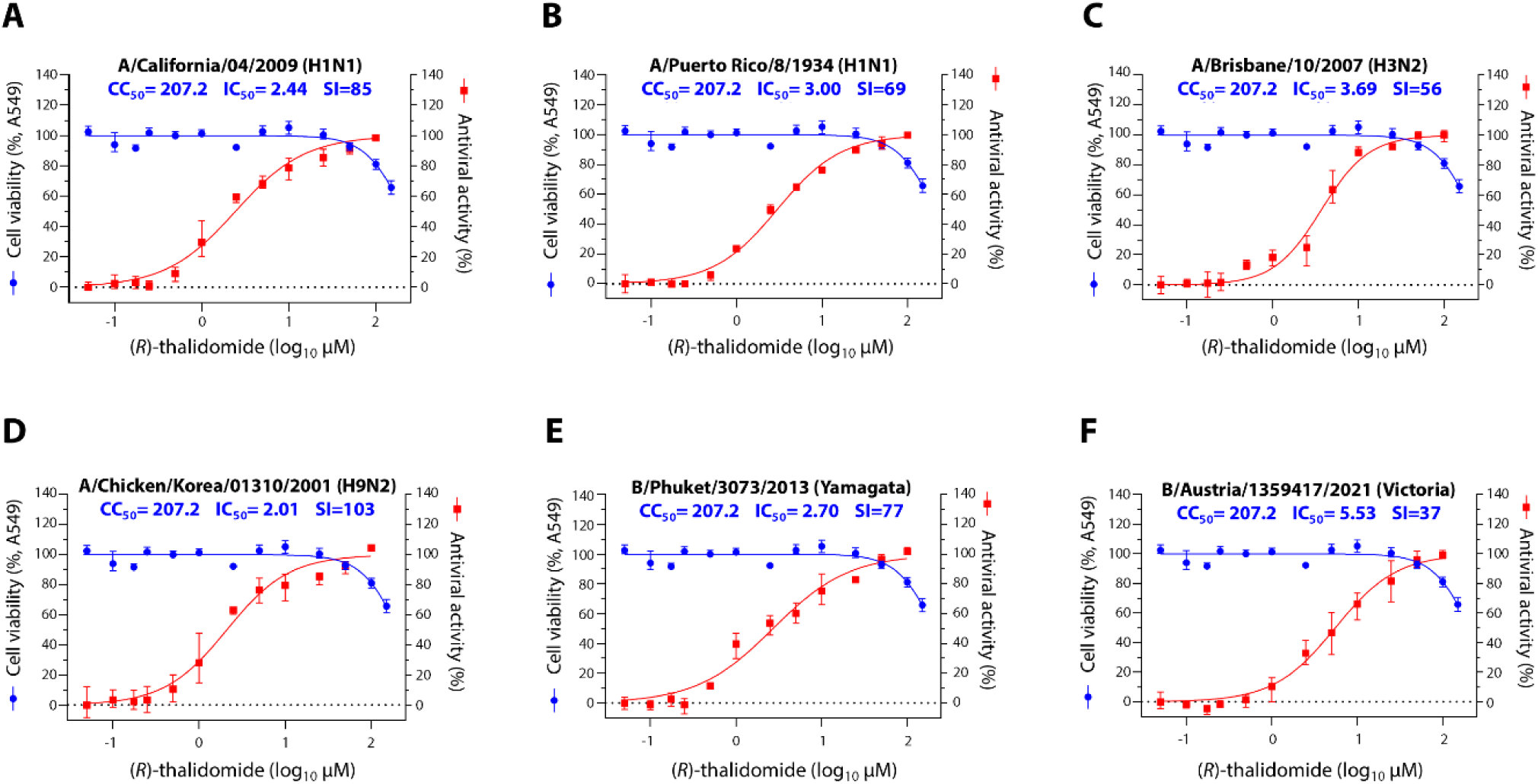
*In vitro* broad-spectrum safety and antiviral effect of (*R*)-thalidomide against different strains of influenza A virus (IAV) and influenza B virus (IBV) (A-F) Dose-response curves displaying the half-maximal inhibitory concentration (IC_50_) and half-maximal cytotoxicity concentration (CC_50_) along with the selective index (SI) of (*R*)-thalidomide against various IAV and IBV strains: A/California/04/2009 (H1N1) pandemic flu (A), A/Puerto Rico/8/1934 (H1N1) seasonal flu (B), A/Brisbane/10/2007 (H3N2) seasonal flu (C), A/Chicken/Korea/01310/2001 (H9N2) low pathogenic (D), and B/Phuket/3073/2013 (Yamagata lineage) seasonal flu (E), and B/Austria/1359417/2021 (Victoria lineage) seasonal flu (F) strains. Viral yield in the cell supernatant was quantified using a cell-culture immunofluorescence assay. Cytotoxicity of (*R*)-thalidomide to each cell line was measured through MTT assay. Data in the graphs are expressed as arithmetic means ± S.D. from four independent experiments. A one-way analysis of variance with Tukey’s correction for multiple comparisons.

**Figure S7.**
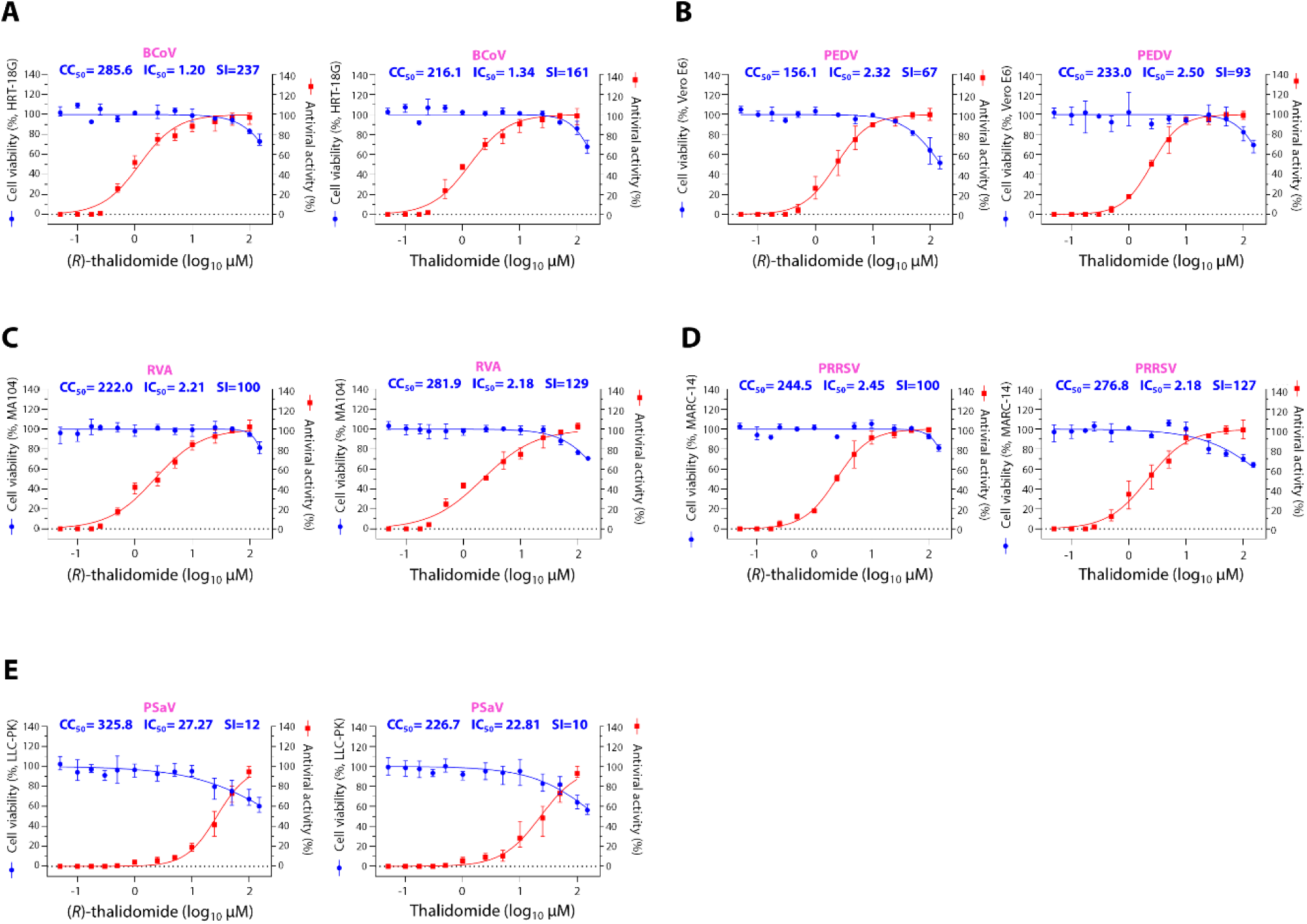
*In vitro* broad-spectrum safety and antiviral effect of (*R*)-thalidomide and thalidomide against different RNA viruses. (A-E) Dose-response curves displaying the half-maximal inhibitory concentration (IC_50_) and half-maximal cytotoxicity concentration (CC_50_) along with the selective index (SI) of (*R*)-thalidomide (left) and thalidomide (right) against different RNA viruses: BCoV KWD strain (A), PEDV KWD strain (B), RVA NCDV strain (C), PRRSV LMY strain (D), and PSaV Cowden strain (E). Viral yield in the cell supernatant was quantified using a cell-culture immunofluorescence assay. Cytotoxicity of (*R*)-thalidomide and thalidomide to each cell line was measured through MTT assay. Data in the graphs are expressed as arithmetic means ± S.D. from four independent experiments. A one-way analysis of variance with Tukey’s correction for multiple comparisons.

**Figure S8.**
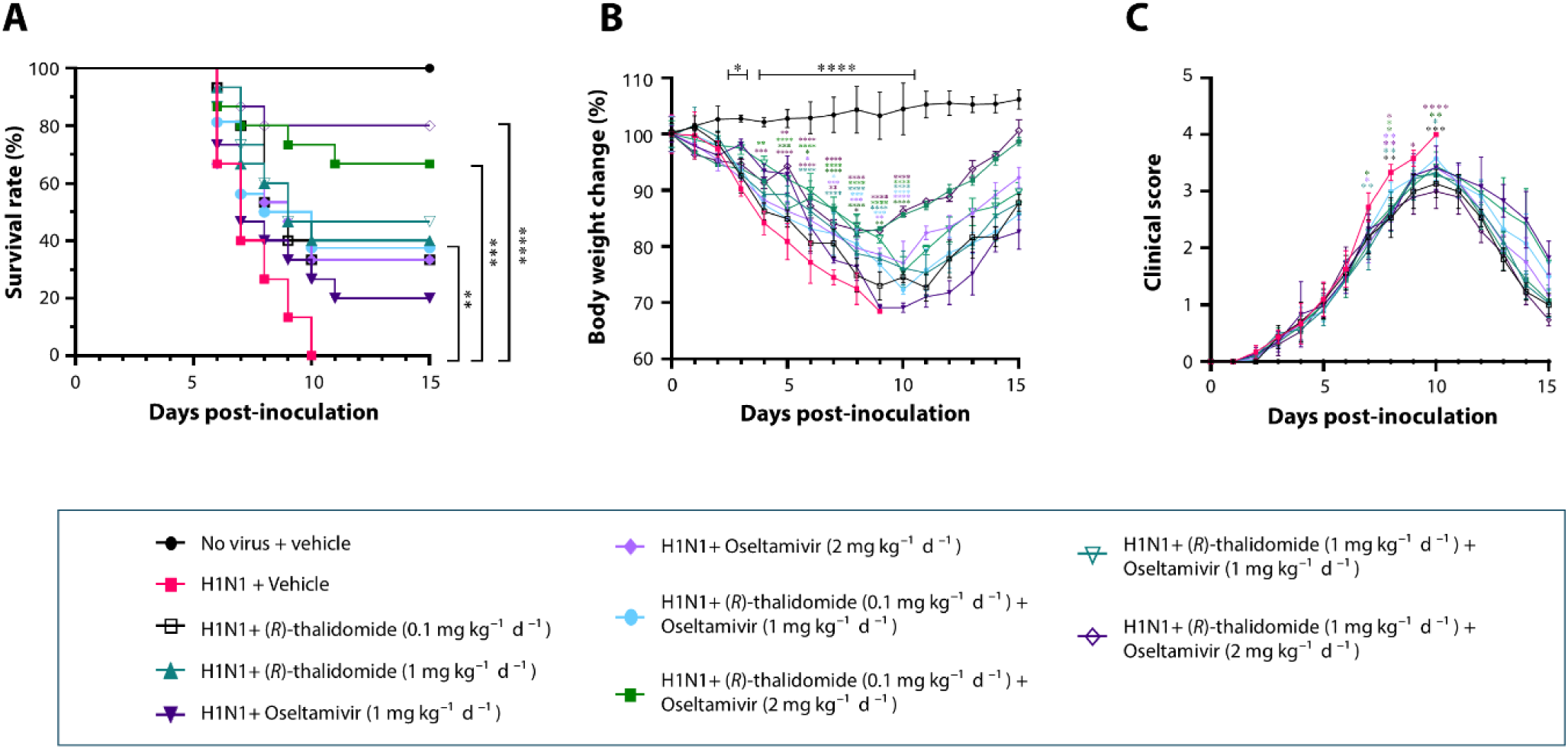
*In vivo* antiviral effects of combination therapy of (*R*)-thalidomide with oseltamivir against influenza A virus (IAV) (A-C) Survival rates (A), daily body weight changes (B), and daily clinical scores (C) of mice administered the (*R*)-thalidomide or oseltamivir, either singly or in combination at different doses for four consecutive days, 1.5 days after challenge with 10^3^ PFU of mouse-adapted IAV PR8 (H1N1) strain. Results are presented as arithmetic means ± S.D. \**P* < 0.05; \*\**P* < 0.01; \*\*\**P* < 0.001, \*\*\*\**p* < 0.00001, a one-way analysis of variance with Tukey’s correction for multiple comparisons.

## Acknowledgments

We thank Professor Ulrich Desselberger at the Department of Medicine, University of Cambridge, for critical reading and discussion, and Dr. Y.B. Baek at the College of Veterinary Medicine, Chonnam National University, for his technical guidance. This work was supported by the National Research Foundation of Korea (NRF) grants funded by the Korean government (MSIT) (RS-2023-00219517 and RS-2024-00339845).

## Author contributions

**Conceptualization:** Thu Ha Nguyen, Dong Ju Lee, Mahmoud Soliman.

**Formal analysis:** Thu Ha Nguyen, Dong Ju Lee, Mahmoud Soliman, Muhammad Sharif.

**Funding acquisition:** Kyoung*-*Oh Cho.

**Investigation:** Myra Hosmillo, Dae-Eun Cheong, Hae-Rang Se, Ian G. Goodfellow.

**Methodology:** Thu Ha Nguyen, Dong Ju Lee, Mahmoud Soliman, Muhammad Sharif, Sunwoo Lee, Don-Kyu Kim, Chul-Seung Park, Hueng-Sik Choi, Tae-Il Jeon, Ian G. Goodfellow, Kyoung*-*Oh Cho.

**Project administration:** Kyoung*-*Oh Cho.

**Resources:** Hyung-Jun Kwon, Seong-Hun Jeong, Sunwoo Lee, Don-Kyu Kim, Chul-Seung Park, Hueng-Sik Choi, Kyoung*-*Oh Cho.

**Supervision:** Tae-Il Jeon, Ian G. Goodfellow, Kyoung*-*Oh Cho

**Validation:** Myra Hosmillo, Dae-Eun Cheong, Hae-Rang Se, Ian G. Goodfellow

**Visualization:** Thu Ha Nguyen, Dong Ju Lee, Mahmoud Soliman, Kyoung*-*Oh Cho

**Writing-original draft:** Thu Ha Nguyen, Dong Ju Lee, Mahmoud Soliman, Sunwoo Lee, Don-Kyu Kim, Chul-Seung Park, Hueng-Sik Choi, Tae-Il Jeon, Ian G. Goodfellow, Kyoung*-*Oh Cho.

**Writing-review & editing:** Thu Ha Nguyen, Dong Ju Lee, Mahmoud Soliman, Sunwoo Lee, Don-Kyu Kim, Chul-Seung Park, Hueng-Sik Choi, Tae-Il Jeon, Ian G. Goodfellow, Kyoung*-*Oh Cho.

